# Assessing the distribution and determinants of vaccine underutilization in the United States

**DOI:** 10.1101/113043

**Authors:** Sandra Goldlust, Elizabeth C. Lee, Murali Haran, Pejman Rohani, Shweta Bansal

**Affiliations:** Department of Biology, Georgetown University, Washington DC, USA; Department of Statistics, Penn State University, University Park PA, USA; Odum School of Ecology and the Department of Infectious Diseases, University of Georgia, Athens GA, USA; Fogarty International Center, National Institutes of Health, Bethesda MD USA

## Abstract

Despite advances in sanitation and immunization, vaccine-preventable diseases remain a significant cause of morbidity and mortality worldwide. In high-income countries such as the United States, coverage rates for vaccination against childhood infections remains high. However, the phenomenon of vaccine hesitancy makes maintenance of herd immunity difficult, impeding global disease eradication efforts. Reaching the ‘last mile’ will require early detection of vaccine hesitancy (driven by philosophical or religious choices), identifying pockets of susceptibility due to underimmunization (driven by vaccine unavailability, costs ineligibility), determining the factors associated with the behavior and developing targeted strategies to ameliorate the concerns. Towards this goal, we harness high-resolution medical claims data to geographically localize vaccine refusal and underimmunization (collectively, ‘underutilization’) in the United States and identify the socio-economic determinants of the behaviors. Our study represents the first large-scale effort for vaccination behavior surveillance and has the potential to aid in the development of targeted public health strategies for optimizing vaccine uptake.

## Introduction

In recent years, vaccine hesitancy (i.e., a desire to delay or refuse vaccination, despite availability of vaccination services) has resurged in the United States, challenging the maintenance of herd immunity.^1^ In December 2014, foreign importation of the measles virus to Disney theme parks in Orange County, California resulted in an outbreak of 111 measles cases, 45% of which were among unvaccinated individuals.^2^ The CDC's National Immunization Survey (NIS) data from 2003 found that 21.8% of parents intentionally delayed vaccine doses, and by 2009 this number had grown to 39.8%.^3^ Effective state legislation and school immunization requirements have kept average vaccination rates above 90% for most required immunizations, reducing the incidence of childhood infections.^4^ However, somewhat counter-intuitively, the drastic reduction in childhood infections in the United States and around the world has also reduced the perception of risk of vaccine-preventable diseases among the public, while heightening concerns over vaccine safety.^3,5,6^

Vaccine hesitancy has been shown to increase the risk of outbreaks of childhood infections, such as measles and pertussis, particularly when it clusters, as it increasingly does in the United States.^7^ While immunization requirements for school entry in the U.S. date back to 1922, a heterogeneous patchwork of vaccination exemption rules and an increasing popularity of private and home schooling has created a mosaic of vaccine coverage across the country. Localized clustering of susceptibility to vaccine-preventable diseases, even in states with high overall mean vaccination rates, increases outbreak risk by degrading herd immunity. Previous small-scale studies have shown that non-medical exemptions for vaccines required for school-entry can vary dramatically across counties within a single state. For example, county-level exemption rates ranged from 1-27% in Washington state in the 2006-2007 school year, despite a state-wide average of 6%.^8^ Similar levels of heterogeneity in vaccine exemption rates have been found in other states including Michigan^9^ and Oregon.^10^ Understanding the spatial distribution of vaccine hesitancy and identifying clusters of underimmunization are critical for determining outbreak risk.

Previous studies have investigated the demographics of intentionally unvaccinated infants and have found them more likely to be Caucasian males of married parents, with greater rates of college education, private school attendance, household income, and household size.^11–13^ Parents engaging in vaccine hesitancy cite personal and religious beliefs as motivating their behavior, including concerns about vaccine safety, doubts regarding the necessity of immunizations for child health, and opposition to school immunization requirements.^3,8,14,15^ The thoroughly debunked claim of an association between the MMR vaccine and autism purported by Andrew Wakefield in a redacted 1998 study has continued to fuel concerns about vaccine safety among parents.^16,17^

Many survey tools have been proposed to assess vaccine sentiment as a proxy for vaccination behavior. A recent survey of vaccine perceptions in 67 countries found reduced vaccine confidence to be associated with high-income countries with strong educational and healthcare systems.^18^ Social media has also been used to assess geographic variability in vaccine sentiment.^19^ Recently, the SAGE Working Group on Vaccine Hesitancy proposed a matrix for classifying the variety of hypothesized determinants, including socioeconomic, religious, demographic, and educational factors.^1^ The Working Group also proposed the following definition for vaccine hesitancy: “Vaccine hesitancy refers to delay in acceptance or refusal of vaccines despite availability of vaccine services. Vaccine hesitancy is complex and context specific, varying across time, place and vaccines. It is influenced by factors such as complacency, convenience and confidence.”^1^ While vaccine hesitancy spans a continuum and can be measured by assessing attitudes and beliefs toward fear of infectious diseases and the vaccines used to prevent them, such attitudes or beliefs do not capture all mechanisms of undervaccination which could also be driven by lack of healthcare access, lack of health insurance, discontinuity of care, competing priorities, social norms, provider recommendations, or vaccination laws.^3^ For example, previous work has found vaccine refusal or intentional delay of vaccination to be positively associated with household income, maternal age, college education, and having married parents, but has also found underimmunization (i.e., not being up-to-date on the full recommended vaccine series) to be negatively associated with these same factors,^11,20^ although these findings have been mixed.^21,22^ Moreover, there is evidence to suggest that concerns about vaccine safety and mistrust in medical advice - both of which have been major focuses of efforts to increase vaccine uptake - may be more characteristic of vaccine refusal or delay as opposed to under-immunization.^11^ Here, we focus on both vaccine hesitancy, as defined by the Working Group, as well as the larger issue of undervaccination.

In the United States, surveillance of vaccine uptake for childhood infections is limited in scope and spatial resolution. Most prior studies have relied on CDC's National Immunization Survey (NIS) or school immunization exemption records to assess vaccine uptake across the United States. While NIS estimates are representative and comparable across states, the survey suffers from small sample sizes (15,000 children), low response rates, and low spatial resolution (state-level).^23–27^ School-based records for immunization and exemptions, on the other hand, provide finer spatial resolution, higher response rates, and vaccine-specific information, but are limited in terms of accuracy (a recent study found 22% of exempt children to in fact be fully vaccinated^28^) and are incomparable across states due to differences in data collection methods and data quality.^29^ A few states have published county-level exemption data, which have been used to study child vaccination trends in California,^30–32^ Michigan,^9^ Washington,^8^ Colorado,^33^,and others.^34^ Surveys have also been conducted to explore the drivers of vaccine hesitancy; however, these studies are typically small-scale and insufficient in-depth exploration of the factors associated with vaccination behavior.^35,36^ The lack of verified tools for assessing and quantifying vaccine underutilization prohibits the development of geographically-targeted and context-specific interventions.^37^ However, electronic health records offer new opportunities to study vaccination behavior using high-volume and high-resolution data with greater clinical accuracy and confirmation.^38^ A limited number of studies have considered the use of ICD-9 codes for assessing vaccination status using claims data from the CDC's Vaccine Safety Datalink (VSD)^39,40^ and Kaiser Permanente managed care organization^41^; however, these studies have been small in scale.

In this study, we leverage fine-grain vaccine underutilization data from a high-coverage medical claims database and combine it with a hierarchical Bayesian modeling approach to estimate local pockets of vaccine underutilization across the United States for the years 2012-2015. We also conduct an ecological analysis of the socio-economic determinants of vaccine underutilization by focusing specifically on vaccine refusal behavior (the rejection of vaccination by choice) versus underimmunization (due to vaccine ineligibility, unavailability or cost). A reliable spatial and temporal understanding of vaccine underutilization, based on both behavioral data and an understanding of the underlying drivers of behavior, could play a critical role in clinical practice and public health decision making.

## Results

We develop a two-level spatial hierarchical Bayesian model to estimate the geographic distribution of vaccine underutilization. Vaccine underutilization is informed by our high-coverage U.S. claims data on cases of vaccine refusal or of underimmunization, as identified by healthcare providers during patient visits. Our model proceeds in two levels: first, we account for imperfect detection of vaccine underutilization cases through medical claims due to spatial variation in healthcare access and insurance rates. Then, based on our spatial estimates of probabilities of detection, we can infer the true abundance and geographic distribution of cases. The model thus allows us to carry out surveillance of vaccine underutilization behavior across the United States. In addition, our model allows us to infer the social, economic, and health policy factors associated with vaccine underutilization, for which we consider vaccine refusal and underimmunization separately.

### Model Fit

The model estimates tend closely to the observed data. Figure 1 shows the model estimates compared to the observed data for each of three outcome measures. The Pearson cross-correlation between the model estimated incidence and the incidence of observed data were R=0.997 (95% CI:0.997-0.997), R=0.986 (95%CI: 0.985-0.987), and R=0.996 (95%CI: 0.996- 0.997), for the model outcomes of vaccine refusal, history of underimmunization, and the combined data, respectively.

**Figure 1.**
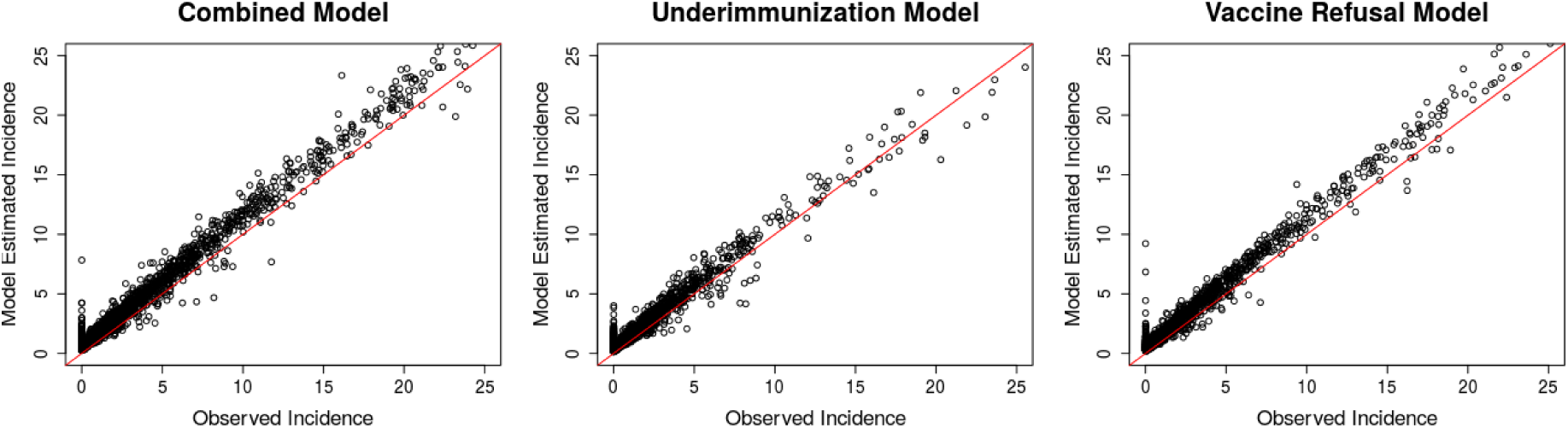
Model fit: Data model estimates of incidence vs. observed incidence (calculated using *μ_i_p* and *y_ij_*, respectively, per 1,000 population under five years of age. Combined (left), underimmunization (middle), vaccine refusal (right). Red line is the equality line (y = x). For clarity, only 2015 data are shown in figure.

### Spatial distribution of vaccine hesitancy and probability of detection

Figure 2 shows our model estimates for vaccine underutilization generated using the combined model. Corresponding model estimates for vaccine refusal and underimmunization alone are shown in Supplemental Figure S2. We observed substantial spatial heterogeneity in vaccine under-utilization and the probability of detection in our database both between and within states.

**Figure 2.**
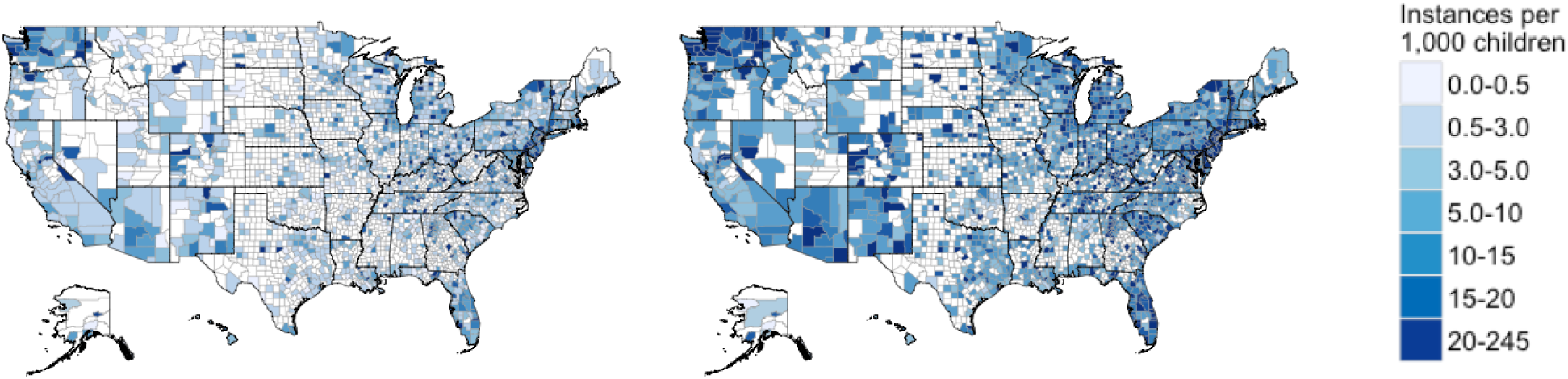
Observed cases and model estimates of vaccine under-utilization from the combined model. Incidences are calculated per 1,000 children under five years of age. Left: Observed combined claims data for 2015. Right: Model estimates of the mean underlying latent incidence of vaccine under-utilization, *μ_i_*, after accounting for detection error. Note: Inconsistently spaced scales. Counties marked NA had no vaccine under-utilization claims over the four years investigated.

**Figure 3.**
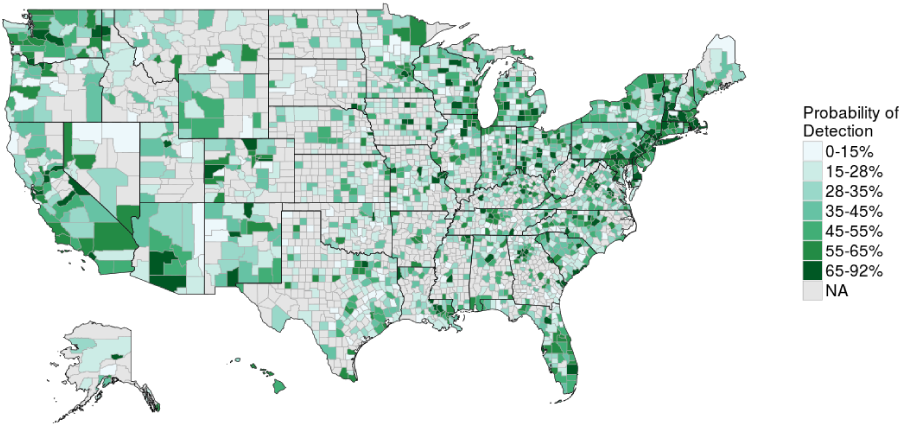
Probability of Detection, *p_i_*,*_j_*, (expressed as percent) in 2015 for the combined model. Missing values shown in grey.

### Predictors of vaccine underutilization and detection

We modeled the two different processes leading to vaccine underutilization (vaccine refusal and underimmunization) separately, to study the determinants of each process (Figure 4). We found that vaccine refusal is positively associated with adult education level and religious adherence, corroborating previous studies.^42,43^ We also found that underimmunization is positively associated with income inequality and negatively associated with percent of population living in the same area one year prior (a measure of continuity of care).^44,45^ However, state laws, state health expenditure, population density, and average family size were not significantly predictive of vaccine refusal or underimmunization, contrary to past studies.^11,18,20,30,34^ Our method also allows us to understand determinants of the measurement of a vaccine underutilization case. We found that access to healthcare (measured through availability of pediatricians) and reporting bias (measured through provider tendency to report a diagnosis on a claim) are both positively associated with both outcome measures (underimmunization and vaccine refusal).

**Figure 4.**
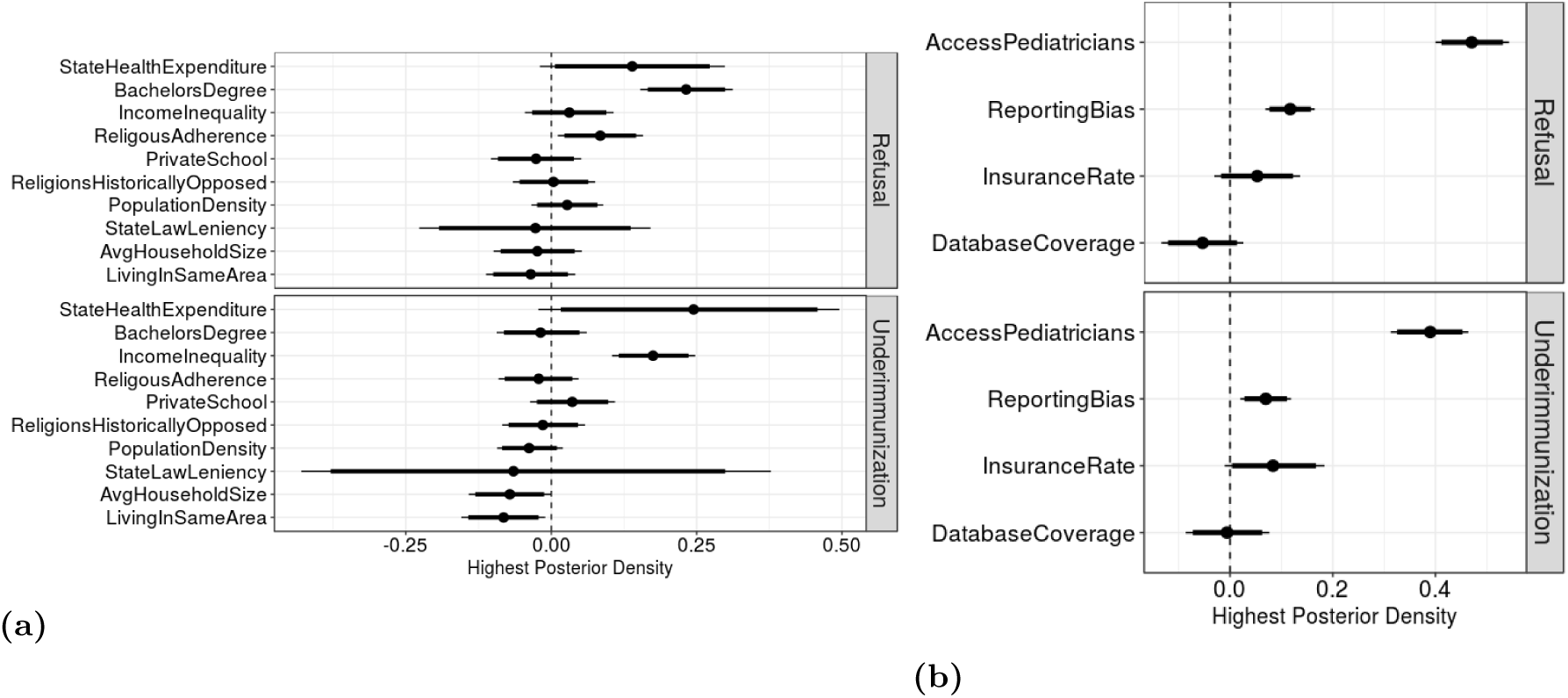
Highest posterior densities (HPD) of the centered and standardized predictors for a) vaccine underutilization and b) probability of detection for vaccine refusal model (top row) and Underimmunization model (bottom row). Thin lines show the 95% HPD region and thick lines show the 90% HPD region. Parameters with intervals crossing the dotted line at HPD = 0 are not significantly associated with the outcome.

### Validation of Results

We conducted model validation for counties within select states with available county-level school vaccine exemption data. (Figure 5) We obtained data on medical and non-medical vaccine exemptions for Kindergartners for the 2014-2015 school year from the California Department of Public Health, Washington State Department of Health, the Texas Department of State Health Services (only non-medical exemption data available), and the New Mexico Department of Health. The Pearson cross-correlations between the modeled latent incidence of vaccine underutilization and the percent of Kindergarteners with a vaccine exemption by county were *R* = 0.53 (*p <* 0.01) for California, R = 0.17 (*p* = 0.055) for Texas, and *R* = 0.40 (*p* = 0.041) for New Mexico.

**Figure 5.**
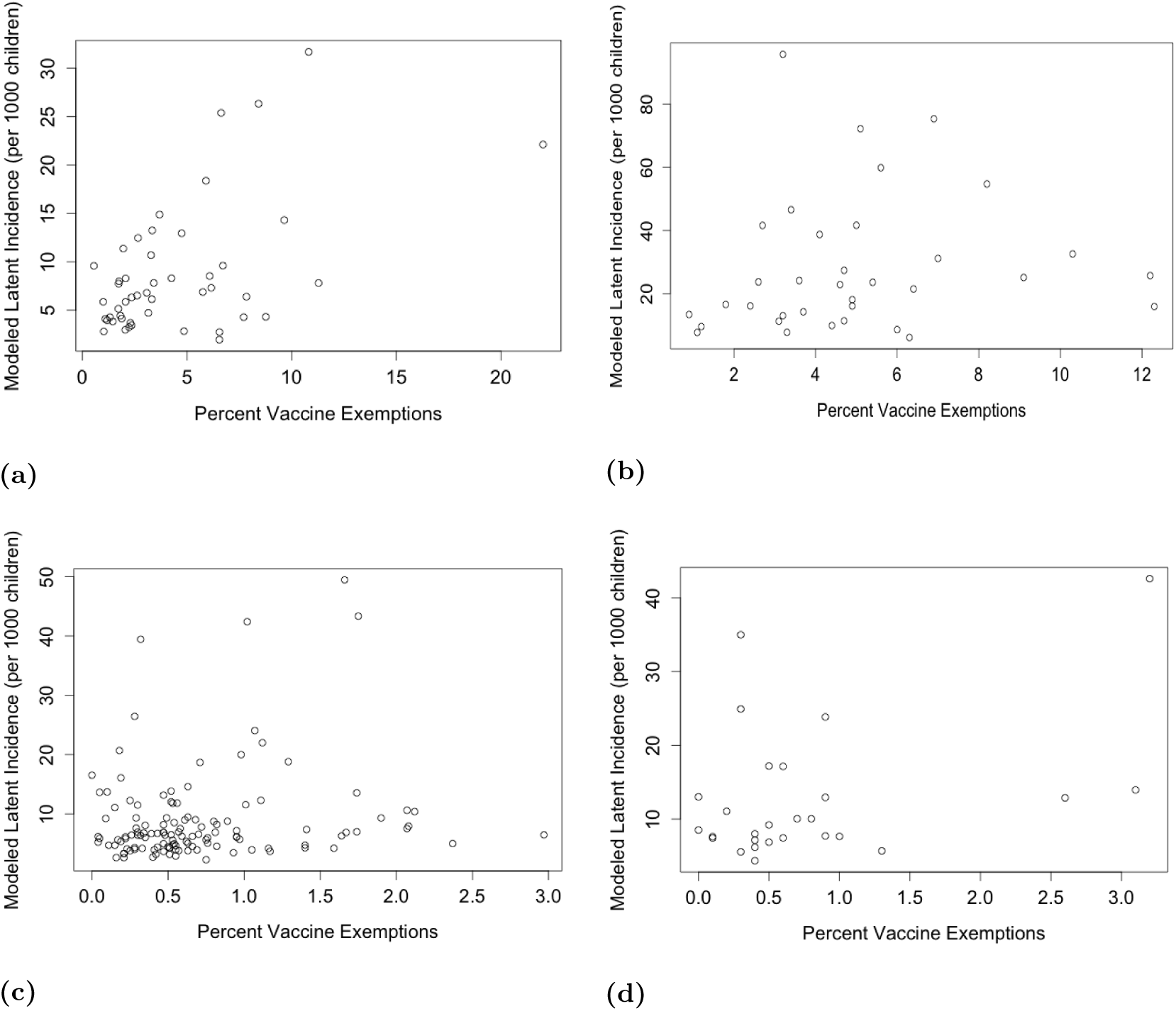
Validation of modeled latent incidence of vaccine underutilization per 1,000 children under 5 years of age compared to available county-level school immunization data on percent of kindergarteners with a vaccine exemptions by county for a) California, b) Washington State c) Texas, and d) New Mexico.

## Discussion

We use a large-scale database of medical claims to create county-level Bayesian maps of the occurrence of vaccine underutilization in the United States for the years 2012-2015. Extending the same models, we quantify the association between various epidemiological factors and vaccination status. To our knowledge, our work is the first to conduct large-scale surveillance of vaccine underutilization across the United States and to investigate the associated behaviors with simultaneously high spatiotemporal resolution and coverage. Our study illustrates the potential of big data to aid in surveillance of vaccine uptake in the United States and study the social determinants of vaccine hesitancy and underimmunization.

Our results help disentangle the process of under-immunization from the process of vaccine refusal. Our observation that underimmunization is positively associated with income inequality suggests that underimmunization may be associated with factors that affect healthcare access. We also observed a negative association between underimmunization and the percent of the population living in the same county one year prior, which may also indicate that continuity of health care plays a significant role in underimmunization. In contrast, we found vaccine hesitancy to be significantly positively associated with the proportion of children attending private schools and rate of religious adherence in the county. These findings suggest may indicate that personal beliefs and choice drive vaccine refusal, while healthcare access and continuity drive underimmunization.

Consideration must be taking to avoid ecological fallacy when interpreting our results. We conduct statistical inference at the county-level, and we therefore are not trying to infer individual factors but rather ecological ones. Previous investigations of disease. distributions has demonstrated sensitivity of statistical inference to spatial scale.^46^ We acknowledge that our medical claims data are not vaccine-specific. Thus, our data may capture vaccine under-utilization for seasonal influenza, human papillomavirus, or other vaccines for which alternative processes drive under-utilization. However, we believe the impact of non-childhood immunizations on our results is limited because we only analyze medical claims submitted for children under five years.

## Materials and Methods

### Medical Claims Data

Monthly reports for vaccine refusal and underimmunization among patients under five years of age were obtained from a database of U.S. medical claims managed by IMS Health. Claims were submitted from both private and government insurance providers, and data were aggregated to U.S. five-digit ZIP codes summarized by year from 2012 through 2015. The reported data cover 359,365 cases of vaccine refusal or underimmunization for children under the age of five across 347,449,121 physician-patient interactions. Vaccine refusal was identified with the International Classification of Diseases, Ninth Revision (ICD-9) code and sub-codes for “vaccination not carried out” (V64.0), thus representing cases when a patient was not immunized due to choice (philosophical refusal or religious reasons). Underimmunization was defined as “personal history of underimmunization status” (V15.83), and thus captures lack of immunization for any reason (vaccine hesitancy, vaccine unavailability, vaccine costs, or vaccine ineligibility). We included counties for which at least one of the four years from 2012-2015 had a non-zero value. In addition, IMS Health provided database metadata on the percentage of reporting physicians and the estimated effective physician coverage by visit volume by three-digit ZIP code (Figure S1). We allocated all ZIP code data to US counties using the address-weighted ZIP Code Crosswalk files provided by the U.S. Department of Housing and Urban Development (HUD) and United States Postal Service (USPS).

### Predictors of vaccine hesitancy

We conducted a literature review to identify hypothesized determinants of vaccine hesitancy, such as income, education, household size, religious group representation, and healthcare policy and expenditures, and searched various publicly-available databases for county or state-level quantifiable measures of these potential drivers. We selected predictor measures initially based on the quality and spatial resolution of available data. We also assessed pairwise correlation and variance inflation factors in order to minimize multi-collinearity before arriving at the final set of predictor measures (Supplemental Table S2). We assumed that there was minimal year-to-year variability in these predictors over the four year period of study and thus obtained one set of spatial data for each predictor to represent the entire study period. All predictor data was centered and standardized for use in the model.

### Predictors of database detection probability

In order to account for variability in measurement bias in our medical claims data, we identified four conditions that would all have to be met in order for an instance of vaccine hesitancy to be captured by our database: 1) seek pediatric health care from a provider 2) be insured; 3) provider use of the claims database 4) the provider reports the vaccine refusal or underimmunization case. Following exploratory analysis and assessing multi-collinearity, we selected four measurement factors representing each of these measurement mechanisms for inclusion in our model (Supplemental TableS1). All predictor data was centered and standardized for use in the model. As we anticipated year-to-year variability in the measurement factors over the four years of our study, we collected four sets of spatial data for each variable in order to account for temporal changes.

### Model Structure

We used a spatial Bayesian hierarchical model to estimate the probability of observing vaccine underutilization in our data, the frequency of vaccine underutilization, and the factors associated with the process of vaccine underutilization. The model structure is similar to an N-Mixture model^47^, which is commonly used for modeling species abundance in ecology. In particular, we separated the processes that describe the probability of observing vaccine underutilization in our data from those that explain the true, latent prevalence of vaccine underutilization in the community. We make a simplifying assumption that the latent prevalence of vaccine underutilization is constant over the duration of our study period, and that our yearly counts of vaccine refusal and underimmunization represent independent replicates of that latent state.

Thus, we modeled three outcomes: the total number of claims submitted to our medical claims database for children under five years of age for 1) vaccine refusal, 2) history of underimmunization, and 3) summation of vaccine refusal and history of underimmunization. For each U.S. county, we assume there exists a latent, true number of instances of cases, and each year in the study period presents an independent opportunity to observe this true state.

We modeled the observed counts of vaccine underutilization (*y_it_*) in county *i* in year *t* with a binomial distribution:

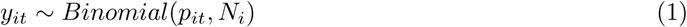

where there is a probability *p_it_* of observing vaccine underutilization in the medical claims among *N_i_*instances in the community (Equation 1). We modeled the probability of observing vaccine underutilization:

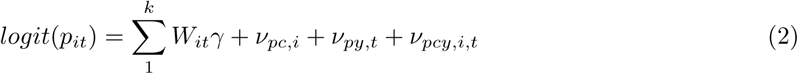

where measurement process predictor *W_it_* is modulated by the estimated coefficient γ, and ν_*pc*,*i*_, ν_*py,t*_, and ν_*pcy,i,t*_ are the group effects for county, year, and county-year, respectively (Equation 2).

In the second level of the hierarchy, we model the latent instances of vaccine underutilization *N_i_*:

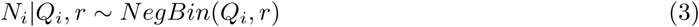

where the negative binomial distribution is parametrized by probability *Q_i_* and size *r* (Equation 3). The mean of *N_i_*, μ_*i*_ is defined:

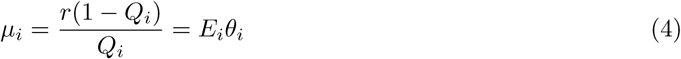

and it can be separated into the expected number of cases *E_i_*, which we calculate by multiplying the country-wide average rate of vaccine under-utilization according to our observed claims data by the population of children under five years of age in county, *i*, and the relative risk θ_*i*_ of vaccine underutilization in county *i* (Equation 4). Finally, the relative risk θ_*i*_ is modeled by:

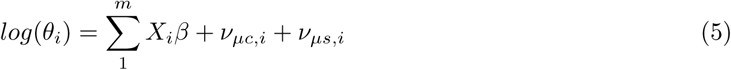

where a single predictor of underutilization *X_i_* is modulated by the estimated coefficient *β* for all *m* predictors, and *ν_μc,i_* and *ν_μs,i_* are county-level and state-level group effects, respectively (Equation 5). Priors and hyperpriors for all *β, γ, r*, and random effects are shown in Supplemental table S4. See Supplemental table S3 for additional definitions of model variables. Estimates of the posterior distributions of model parameters were generated via Markov chain Monte Carlo (MCMC) methods using JAGS and the runjags package in R.^48–50^ Parameter convergence was evaluated by assessing traceplots, the Gelman-Rubin convergence diagnostic, autocorrelation, Monte Carlo standard error, and effective sample size.^51,52^ We evaluated model residuals and compared modeled and observed outcomes in order to assess model fit.

### Validation

Results validation was performed by comparing our model estimates to county-level data from annual school vaccine assessments reports on the rates of vaccine exemptions for Kindergarteners. We selected states for validation based on the quality and availability of their county-level vaccine exemption data. Due to state differences in the quality and collection methods of vaccine exemption data, we validated our results separately for each selected state.

## Supplemental Information

### Predictor data of Detection Probability

In Table S1, we list the predictors used to predict detection probabilities. These predictors vary annually and spatially (by county). To control for variation in physician reporting practices, we compared use of low birth weight ICD-9 codes (subcodes of V21.3, 765.0, and 765.1) in our claims data with county-level birth weight data obtained from the CDC National Vital Statistics system.

**Table S1.** Predictor Data of Detection Probability.

| Label | Measure | Scale | Year(s) | Source |
| --- | --- | --- | --- | --- |
| AccessPediatricians | Pediatricians per population | county | 2011-2014 | AHRF |
| DatabaseCoverage | Database Coverage | county | 2012-2015 | IMS Health |
| InsuranceRate | Health insurance rate | county | 2011-2014 | SAHIE |
| ReportingBias | ICD-9 reporting rate | county | 2012-2015 | CDC Natality;<br>IMS Health |

### Predictor data of Vaccine hesitancy/underimmunization

In Table S2, we list the predictors used for the vaccine refusal or under-immunization process.

**Table S2.** Predictor Data of Vaccine Under-utilization

| Label | Measure | Scale | Year(s) | Source |
| --- | --- | --- | --- | --- |
| StateHealthExpenditure | Healthcare Expenditure | state | 2009 | CMS |
| BachelorsDegree | Percent with bachelors degree | county | 2011-2015 | ACS |
| IncomeInequality | Income Inequality: Ratio of household income of 80th percentile to that of 20th percentile | county | 2010-2014 | ACS |
| ReligiousAdherence | Percent religious adherents | county | 2010 | RCMS |
| PrivateSchool | Percent private school enrollment | county | 2011-2015 | ACS |
| ReligionsHistoricallyOpposed | Per capita congregations of religions historically opposed to vaccination | county | 2010 | RCMS |
| PopulationDensity | Population density | county | 2010 | ACS |
| StateLawLeniency | Leniency of state vaccination laws | state | 2015 | CDC |
| AvgHouseholdSize | Average household size | county | 2011-2015 | ACS |
| LivingInSameArea | Percent of population living in the same county one year prior | county | 2010-2014 | AHRF |

Measures excluded after variable selection: Child poverty rate (SAIPE), percent of children living in single-parent households (ACS), high school graduation rate (ACS), median income (SAIPE), hospital utilization (AHRF). ACS = American Community Survey, CDC = Centers for Disease Control and Prevention, CMS = Centers for Medicare and Medicaid Services, AHRF = Area Health Resource Files. SAIPE = Small Area Income and Poverty Estimates, RCMS = U.S. Religion Census: Religious Congregations and Membership Study

**Table S3.** Model Variables.

| Variable | Definition |
| --- | --- |
| $N_i$ | Latent state of county, $i$ |
| $y_{i,t}$ | Observed response at county, $i$, in year, $t$ |
| $\mu_i$ | Mean of the distribution of latent response of county, $i$ |
| $E_i$ | Expected response of county (offset), $i$, calculated directly |
| $p_{i,t}$ | Probability of detection at county, $i$, in year, $t$ |
| $Q_i$ | Negative binomial probability parameter of county, $i$ |
| $r$ | Negative binomial size parameter |
| $\beta$ | process model coefficients |
| $x_i$ | county-level process drivers |
| $\gamma_1$ | detection model coefficients |
| $w_{i,t}$ | county-year-level detection drivers |
| $\nu_{\mu c,i}$ | Unstructured county random effects on $\mu_i$ |
| $\nu_{\mu s,i}$ | Unstructured state random effects on $\mu_i$ |
| $\nu_{pc,i}$ | Unstructured county random effects on $p_{i,t}$ |
| $\nu_{py,t}$ | Unstructured year random effects on $p_{i,t}$ |
| $\nu_{pcy,i,t}$ | Unstructured county-year random effects on $p_{i,t}$ |
| $i, t$ | county, year, indicators, respectively |
| $\sigma_{\mu c}$ | standard deviation of random effect $\nu_{\mu,i}$ |
| $\sigma_{\mu s}$ | standard deviation of random effect $\nu_{\mu,i}$ |
| $\sigma_{pc}$ | standard deviation of random effect $\nu_{\mu,i}$ |
| $\sigma_{py}$ | standard deviation of random effect $\nu_{py,t}$ |
| $\sigma_{pcy}$ | standard deviation of random effect $\nu_{pcy,i,t}$ |

Description of data, *y_i,t_*: county(*i*)/year(*t*)-level counts of patient claims for history of under-immunization for the year.

**Table S4.**
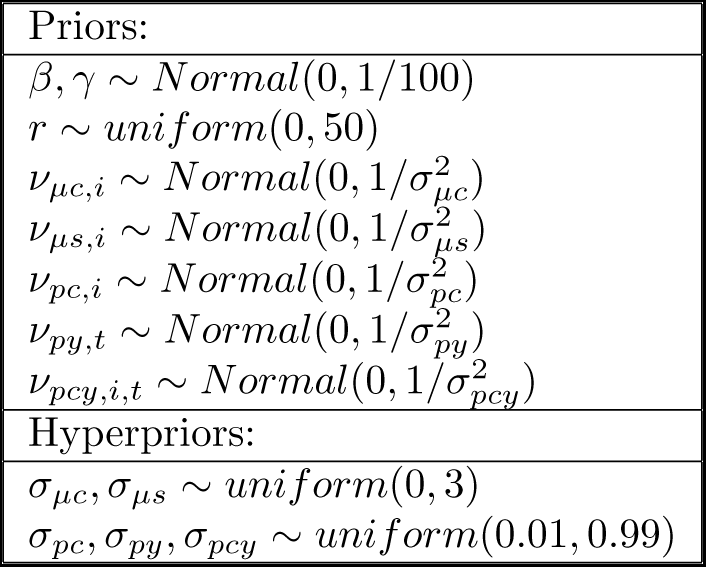
Priors and Hyperpriors.

We initialized parameter coefficients β and γ using each with a random draw from a uniform distribution from −1 to 1. Standard deviation hyperpriors for the group-level effects and *r* were initialized each with a random draw from a uniform distribution 0 to 1, and 1 to 20, respectively. *N_i_* was initialized for each i county as the maximum observation, *y_i,t_* over *t* years, plus a random draw from the Poisson distribution with mean 10.

**Figure S1.**
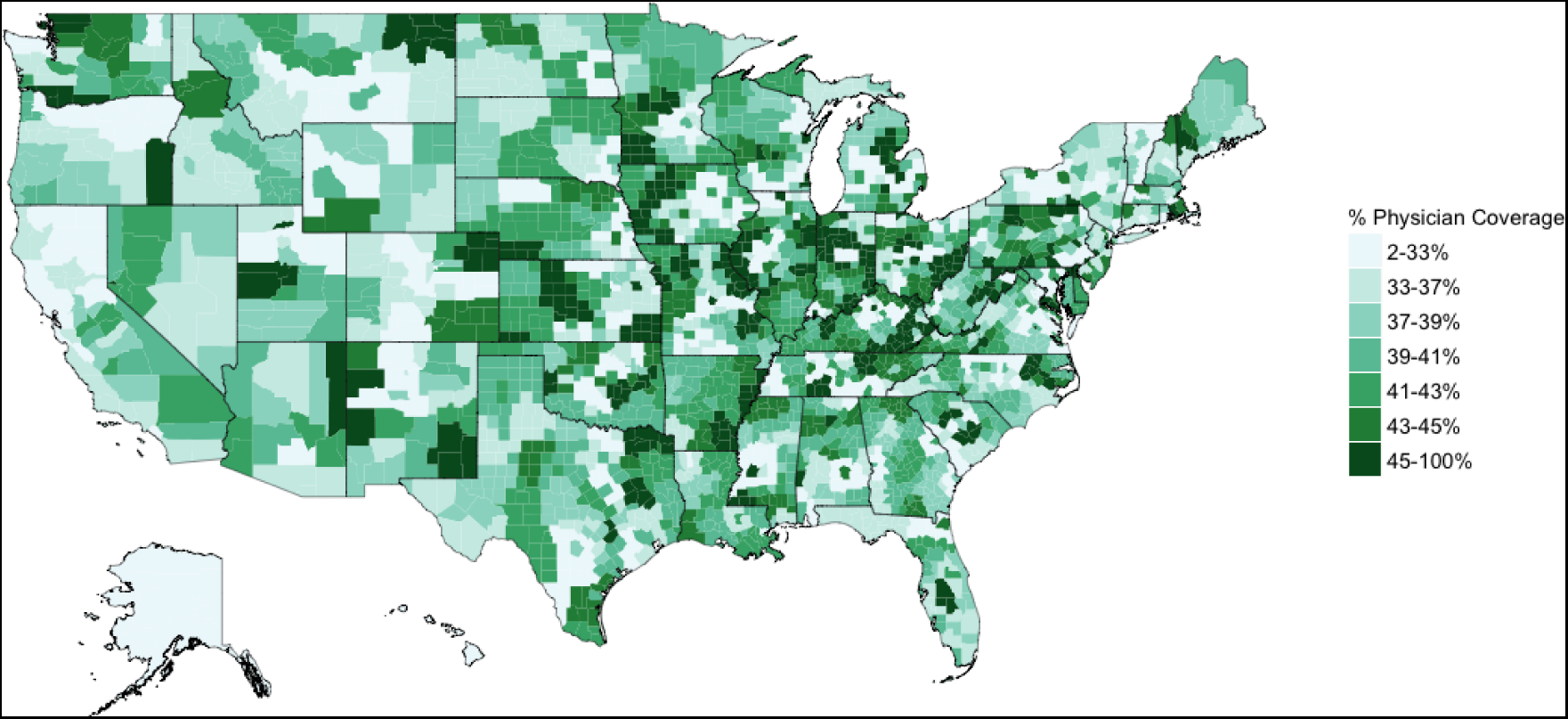
Database physician coverage, 2015. Percent of AMA licensed physicians reporting to our medical claims database by three-digit zip-code and adjusted for estimated visit capture (our database does not always capture 100% of a physician's visits). Our claims database additionally includes reporting from other healthcare providers (nurses, physicians assistants, etc.); however, in order to compare representation to the AMA universe, only AMA physicians are shown here.

**Figure S2.**
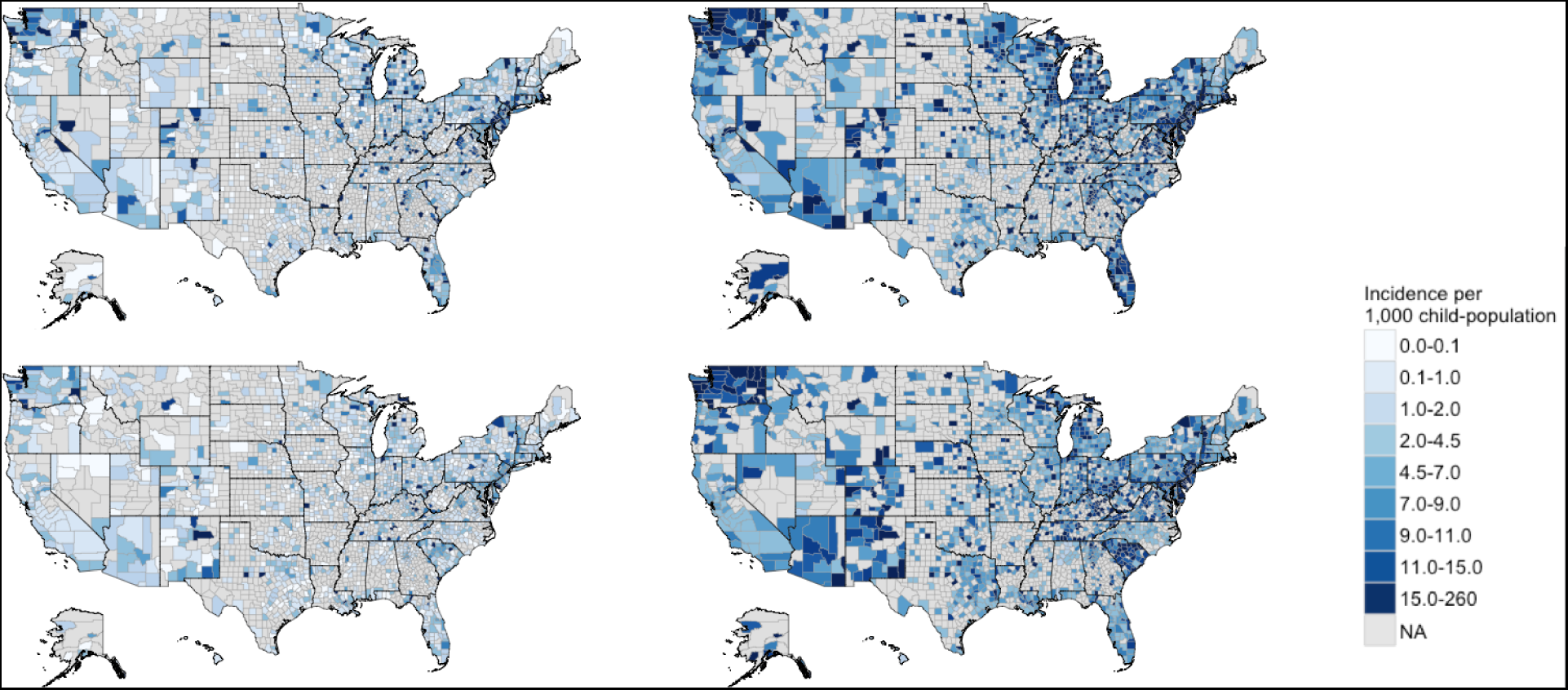
2015 Observed cases (left) and model estimates (right) of vaccine under-utilization from the vaccine refusal model (top row) and underimmunization model (bottom row). Incidences are calculated per 1,000 children under five years of age. Left: Observed refusal claims data for 2015. Right: Model estimates of the mean underlying latent incidence of vaccine under-utilization, μ_*i*_, after accounting for detection error. Note: Inconsistently spaced scales. Counties marked NA had no vaccine under-utilization claims over the four years investigated.

